# Assessment of DNA methylation patterns in the bone and cartilage of a nonhuman primate model of osteoarthritis

**DOI:** 10.1101/184879

**Authors:** Genevieve Housman, Lorena M. Havill, Ellen E. Quillen, Anthony G. Comuzzie, Anne C. Stone

**Affiliations:** School of Human Evolution and Social Change, Arizona State University, Tempe, AZ, USA.; Center for Evolution and Medicine, Arizona State University, Tempe, AZ, USA.; Southwest National Primate Research Center, Texas Biomedical Research Institute, San Antonio, TX, USA.; Department of Genetics, Texas Biomedical Research Institute, San Antonio, TX, USA.

**Author notes:** Author Notes: GH is currently affiliated with the University of Chicago, and EEQ is currently affiliated with Wake Forest University School of Medicine. Corresponding author Genevieve Housman, Section of Genetic Medicine, University of Chicago, 920 East 58th Street, CLSC 317, Chicago, IL 60637, USA. **Funding:** NIH Grant P01 HL028972 to A.G.C; Max and Minnie Tomerlin Voelcker Foundation to L.M.H.; William and Ella Owens Foundation for Biomedical Research to L.M.H.; SNPRC Internship Funds to G.H.; ASU Chapter of Sigma Xi Grant-in-Aid of Research to G.H.

**Keywords:** osteoarthritis, animal model, articular cartilage, bone, DNA methylation

## Abstract

**Objective:** Osteoarthritis (OA) impacts humans and several other animals. Thus, the mechanisms underlying this disorder, such as specific skeletal tissue DNA methylation patterns, may be evolutionary conserved. However, associations between methylation and OA have not been readily studied in nonhuman animals. Baboons serve as important models of disease and develop OA at rates similar to those in humans. Therefore, this study investigated the associations between methylation and OA in baboons to advance the evolutionary understanding of OA.

**Design:** Trabecular bone and cartilage was collected from the medial condyles of adult female baboon femora, five with and five without knee OA. The Infinium HumanMethylation450 BeadChip (450K array) was used to identify DNA methylation patterns in these tissues.

**Results:** Approximately 44% of the 450K array probes reliably align to the baboon genome, contain a CpG site of interest, and maintain a wide distribution throughout the genome. Of the two filtering methods tested, both identified significantly differentially methylated positions (DMPs) between healthy and OA individuals in cartilage tissues, and some of these patterns overlap with those previously identified in humans. Conversely, no DMPs were found between tissue types or between disease states in bone tissues.

**Conclusions:** Overall, the 450K array can be used to measure genome-wide DNA methylation in baboon tissues and identify significant associations with complex traits. The results of this study indicate that some DNA methylation patterns associated with OA are evolutionarily conserved, while others are not. This warrants further investigation in a larger and more phylogenetically diverse sample set.

## Introduction

Osteoarthritis (OA) is a complex degenerative joint disease, and OA of the knee is one of the leading causes of disability across the globe.^1^ Thus, research endeavors to describe the molecular mechanisms that contribute to this disorder are underway. Both genetic and environmental factors have some effect,^2,3^ as well as epigenetic factors which bridge the gap between genetics and the environment. In particular, DNA methylation which regulates gene expression, is thought to play an influential role in the development of degenerative skeletal disorders like OA.^4–10^ Animal models, such as mice, rats, rabbits, guinea pigs, dogs, sheep, goats, and horses, have been essential in discerning some of the processes inherent to OA development.^11,12^ Nevertheless, all of these animals are limited in their abilities to fully inform our understanding of human OA, so the search to find a gold standard animal model for OA is still ongoing.^13^ Lastly, although variation in skeletal tissue DNA methylation patterns are thought to be involved in the development and progression of OA^4–10^ this epigenetic mechanism has not been readily studied in animal models. In part, this may be due to the compatibility constraints of DNA methylation assays that are designed specifically for humans, as well as the limited efforts to optimize these assays for nonhuman animals.^14,15^

Nonhuman primates can serve as important models of disease for humans because they are phylogenetically close to humans. Baboons (*Papio sp*.) are a particularly good model of disease, especially OA,^16^ as they naturally develop OA at rates similar to those observed in humans.^3,16^ Additionally, because of their evolutionary proximity to humans, further investigation of the molecular processes innate to OA development and progression in baboons as compared to these mechanisms in humans will advance the evolutionary understanding of this disease. Lastly, the relative genetic conservation between baboons and humans makes the optimization and use of standardized DNA methylation assays possible. Specifically, the Infinium HumanMethylation450 BeadChip (450K array), which is a cost-efficient application for assessing genome-wide DNA methylation patterns in humans, has been successfully used for some nonhuman primate species. These and other nonhuman primate DNA methylation studies have primarily used DNA extracted from blood or other soft tissues.^14,15,17–20^ However, this technique has not yet been used to study DNA methylation variation in baboon skeletal tissues or how it relates to the development of OA in a nonhuman primate species.

For this study, we used the 450K array to identify DNA methylation patterns in femur bone and cartilage of age-matched female baboons, five with and five without knee OA. This was done to validate that the 450K array can be used for nonhuman primate skeletal tissue DNA extracts. Additionally, this study was performed to determine if DNA methylation variation is associated with OA in baboons and in a manner similar to that observed in humans.

## Methods

### Ethics Statement

Nonhuman primate tissue samples included were opportunistically collected at routine necropsy of these animals. No animals were sacrificed for this study, and no living animals were used in this study.

### Samples

Baboon (*Papio sp*.) samples come from captive colonies at the Southwest National Primate Research Center in Texas. These samples are ideal because many environmental factors that influence skeletal development and maintenance (e.g., diet and exposure to sunlight which influences vitamin D production) are controlled and consistent across individuals. Femora were opportunistically collected at routine necropsy of these animals and stored in -20°C freezers at the Texas Biomedical Research Institute after dissection. These preparation and storage conditions ensured the preservation of skeletal DNA methylation patterns. Samples include skeletally healthy adult baboons (n=5) and adult baboons with severe OA (n=5). Age ranges are comparable between each group (19.30±1.70 years and 19.24±1.73 years, respectively), and only females were included in this study.

### Assessment of Osteoarthritis

Classification of adult baboons as having healthy or OA knees was determined through visual examination of the distal femora and macroscopic inspection of the distal articular surface cartilage. Each specimen was assigned an OA severity score. Briefly, Grade 1 is unaffected, Grade 2 is mild OA as indicated by cartilage fibrillation, Grade 3 is moderate OA as indicated by cartilage lesions, and Grade 4 is advanced OA as indicated by eburnation.^3^ From this, binary classifications were made such that all healthy adult baboons have 100% Grade 1 on one or both distal femora, and all OA adult baboons have a variable percentage of Grades 3 or 4 on one or both distal femora (Figure 1).

**Figure 1.**
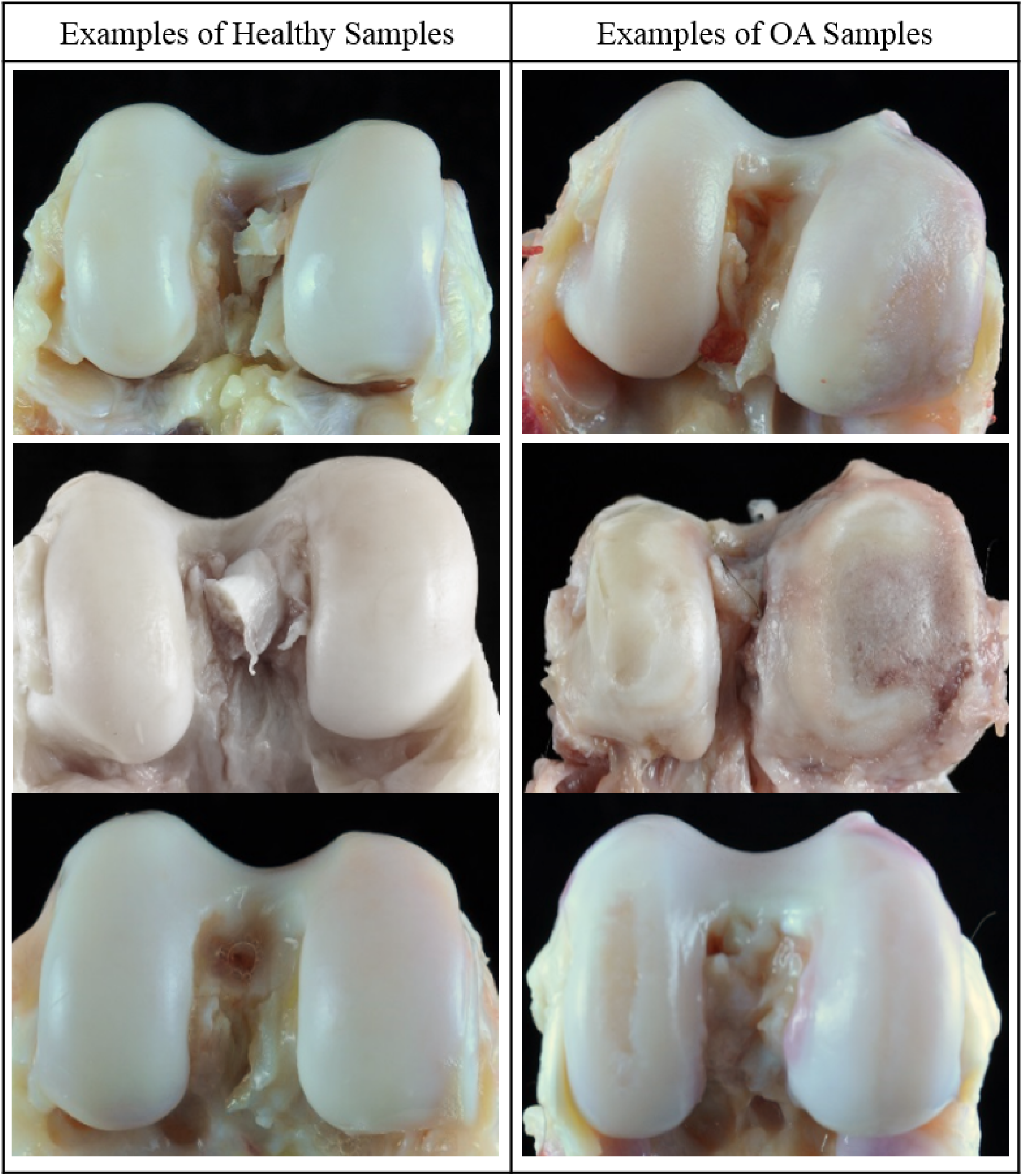
Representative examples of baboon knees (distal femora) that are healthy or have OA.

### DNA Extraction

From the distal femoral condyles, cartilage scrapings were collected using scalpels and processed with a homogenizer, and trabecular bone samples from the interior of the condyle were obtained using a Dremel and pulverized into bone dust using a BioPulverizer. Tissues were collected from this region because OA legions in both humans and baboons generally appear first on the cartilage of the medial condyle. Both tissues are included in this project because human skeletal epigenetic studies are based on trabecular bone and cartilage, and it is important to standardize tissue type for comparative purposes. DNA was extracted from these processed tissues using a phenol-chloroform protocol optimized for skeletal tissues^21^ and quantified using both Nanodrop and Qubit machines (Table S1).

### Genome-Wide DNA Methylation Profiling

Genome-wide DNA methylation was assessed using Infinium HumanMethylation450 BeadChips (450K array). These arrays analyze the methylation status of over 485,000 sites throughout the genome, covering 99% of RefSeq genes and 96% of the UCSC-defined CpG islands and their flanking regions. For each sample, approximately 500ng of genomic DNA (Table S1) was bisulfite converted using the EZ DNA Methylation™ Gold Kit, according to the manufacturer’s instructions (Zymo Research), with modifications described in the Infinium Methylation Assay Protocol. Following manufacturer guidelines (Illumina), this processed DNA was then whole-genome amplified, enzymatically fragmented, hybridized to the arrays, and imaged using the Illumina iScan system. The array data discussed in this publication have been deposited in NCBI’s Gene Expression Omnibus and are accessible through GEO Series accession number GSE101733.

### Processing of Methylation Data

Raw fluorescent data were normalized to account for the noise inherent within and between the arrays themselves. Specifically, we performed a normal-exponential out-of-band (Noob) background correction method with dye-bias normalization^22^ to adjust for background fluorescence and dye-based biases and followed this with a between-array normalization method (functional normalization)^23^ which removes unwanted variation by regressing out variability explained by the control probes present on the array as implemented in the minfi v1.20.2 package in R v3.3.1^24^ which is part of the Bioconductor project.^25^ This method has been found to outperform other existing approaches for studies that compare conditions with known large-scale differences,^23^ such as those assessed in this study.

After normalization, methylation values (β values) for each site were calculated as the ratio of methylated probe signal intensity to the sum of both methylated and unmethylated probe signal intensities. These β values range from 0 to 1 and represent the average methylation levels at each site across the entire population of cells from which DNA was extracted (0 = completely unmethylated sites, 1 = fully methylated sites). Every β value in the Infinium platform is accompanied by a detection p-value, and those with failed detection levels (p-value > 0.05) in greater than 10% of samples were removed from downstream analyses. Because β values have high heteroscedasticity, they are not statistically valid for use in differential methylation analyses.^26^ Thus, M values were calculated as the log transformed ratio of methylated signal to unmethylated signal and used in the statistical analyses described below.

The probes on the arrays were designed to hybridize specifically with human DNA, so our use of nonhuman primate DNA required that probes non-specific in the baboon genome, which could produce biased methylation measurements, be computationally filtered out and excluded from downstream analyses. This was accomplished using two different methods modified from ^14,15^. For both methods, we used blastn^27^ to map the 485,512 50bp probes onto the *Papio anubis* genome (Assembly: Panu_2.0, Accession: GCF_000264685.2) using an e-value threshold of e^-10^. We retained probes that successfully mapped to the baboon genome, had only one unique BLAST hit, and targeted CpG sites (Supplemental Probe Annotation File). These 213,858 probes that aligned to the baboon genome were then further filtered using one of two methods.

The first method used criteria based on sequence alignment^14^, and the second method used criteria based on gene symbol similarities^15^. For the first method (alignment filter), we only retained probes that had 0 mismatches in the 5bp closest to and including the CpG site and that had 0-2 mismatches in the 45bp not including the CpG site.^14^ This more stringent sequence similarity filtering retained 133,264 probes. For the second method (gene symbol filter), we identified the closest baboon gene to each probe alignment site and checked for corresponding gene name matches between humans and baboons.^15^ For baboons, this information was obtained from GFF and Ensembl BioMart data. Only those probes with partial or complete gene matches were retained. This more lenient filtering retained 130,307 probes. Additionally, cross-reactive probes,^28^ probes containing SNPs at the CpG site, probes detecting SNP information, probes detecting methylation at non-CpG sites, and probes targeting sites within the sex chromosomes were removed using the minfi v1.20.2 package in R v3.3.1.^24^ This filtering retained a final set of 120,305 probes for the alignment filter criteria and 112,760 probes for the gene symbol criteria (Figure 2).

**Figure 2.**
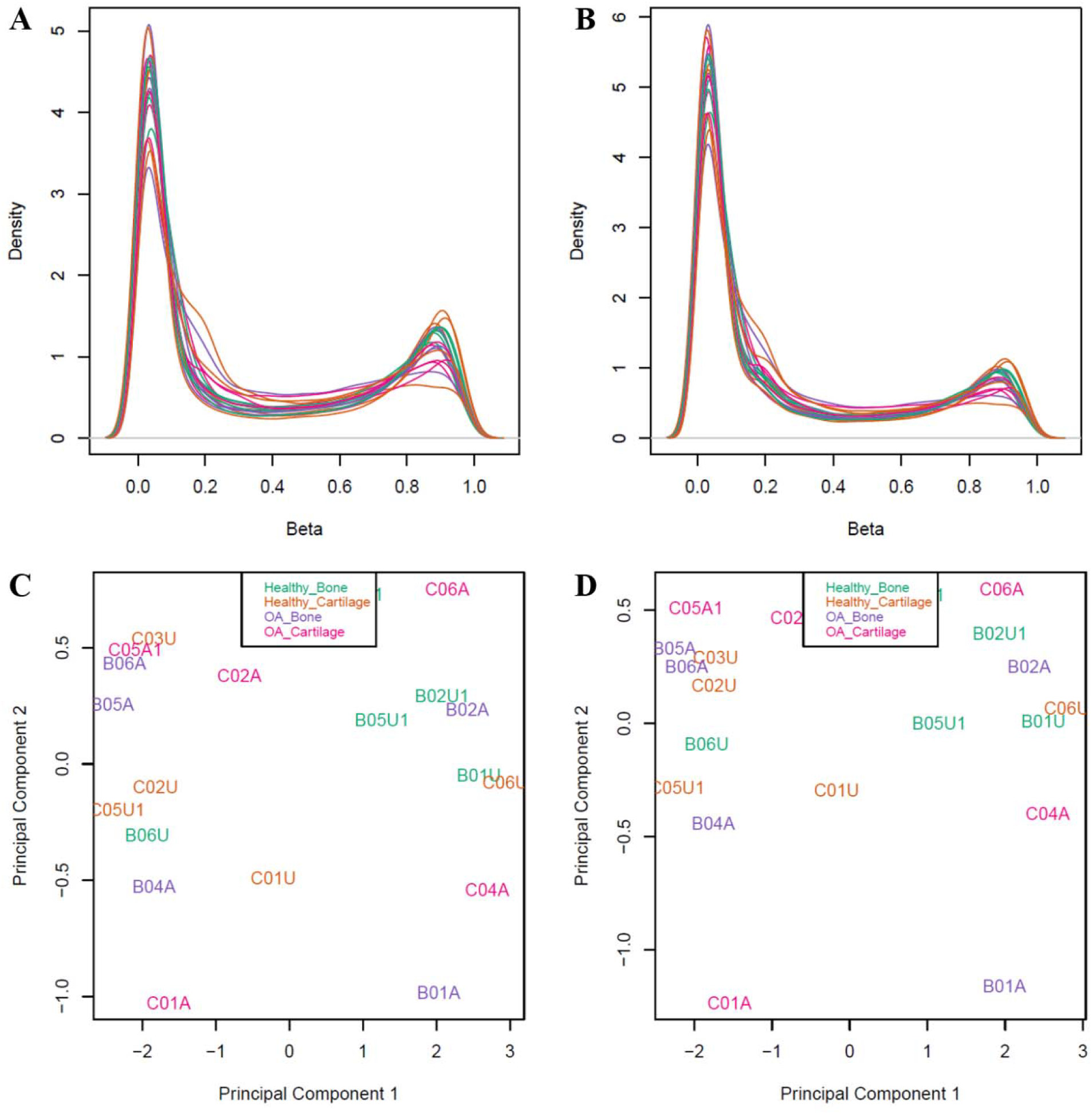
**A** and **B**, Density plots of β values after normalization and probe filtering using the alignment filter criteria **(A)** or the gene symbol filter criteria **(B)**. **C** and **D**, Multidimensional scaling plots showing the first two principle components that describe genome-wide methylation variation after normalization and filtering using the alignment filter criteria **(C)** or the gene symbol filter criteria **(D)**. Each point represents one sample that is either from healthy bone, healthy cartilage, OA bone, or OA cartilage. In the multidimensional scaling plots, these categories do not form distinct clusters.

### Statistical Analysis of Differential Methylation

To identify sites that were significantly differentially methylated across comparative groups, we designed and tested generalized linear mixed models (GLMMs) which related the variables of interest to the DNA methylation patterns for each site, while accounting for latent variables.^29^ Sites found to have significant associations were classified as significant differentially methylated positions (DMPs). Specifically, a GLMM was used to estimate differences in methylation levels for each of the following contrasts: (1) between bone and cartilage in OA baboons, (2) between bone and cartilage in healthy baboons, (3) between OA and healthy baboon bone, (4) between OA and healthy baboon cartilage, (5) among all four combinations of tissue type and disease state (healthy bone vs. healthy cartilage vs. OA bone vs. OA cartilage).

Additional variables included in this GLMM were unknown latent variables calculated using the iteratively re-weighted least squares approach in the sva v3.22.0 package in R v3.3.1.^30,31^ The four latent variables estimated were included to help mitigate any unknown batch and cell heterogeneity effects on methylation variation at each site. No predefined batch effects for the arrays were included because these did not appear to have large effects on the data (Figure S1). Alternative methods to account for cell heterogeneity exist, but they are specific to whole blood,^30,32^ require reference epigenetic data, or are reference free methods^33^ that are comparable to the sva method.^34^ Out of the known cell types in skeletal tissues, only chondrocytes and osteoblasts have reference epigenomes available on the International Human Epigenomics Consortium, and these are only for humans, not nonhuman primates. Thus, because no standard method is available to correct for the heterogenous cell structure in nonhuman primate skeletal tissue, we chose the described sva method.

The GLMM design matrix was fit to the M value array data by generalized least squares using the limma v3.30.13 package in R v3.3.1,^25,35^ and the estimated coefficients and standard errors for the defined tissue type and disease status contrasts were computed. Lastly, for each coefficient, an empirical Bayes approach^36^ was used to compute moderated t-statistics, log-odds ratios of differential methylation, and associated p-values adjusted for multiple testing.^37^ Significant DMPs for the effect of tissue type and disease status contrasts were defined as those having log fold changes in M values corresponding to an adjusted p-value of less than 0.05.

## Results

The aim of this study was to use the 450K array to identify DNA methylation patterns in femur bone and cartilage of age-matched female baboons, five with and five without knee OA. In order to do this, we first assessed the effectiveness of the 450K array in identifying DNA methylation patterns in baboon DNA and of different probe filtering methods.

### Alignment of 450K Array Probes with the Baboon Genome

Probes from the 450K array were aligned to the baboon genome (Supplemental Probe Annotation File).^14, 15^ Out of the 485,512 50bp probes on the array, 213,858 probes (44%) map to the baboon genome with e-values less than e^-10^, have only unique BLAST hits, and target a CpG site ( Figure 3A). Out of these reliably mapped probes, 133,264 probes (62%) were retained after the alignment filter criteria (Figure 3B), while 130,307 probes (61%) were retained after the gene symbol filter criteria (Figure 3C). 83,142 probes overlapped between both filtering methods (62% for the alignment filter criteria and 64% for the gene symbol filter criteria, Figure S2). Probes that reliably mapped to the baboon genome, that met the alignment filter criteria, or that met the gene symbol criteria covered approximately 18,800 genes with an average coverage of nine, six, or eight probes per gene, respectively (Table S2). Additionally, the retained probes covered a range of locations with respect to genes and CpG islands (Table S2), indicating that these filtered probes maintain a wide distribution throughout the genome.

**Figure 3.**
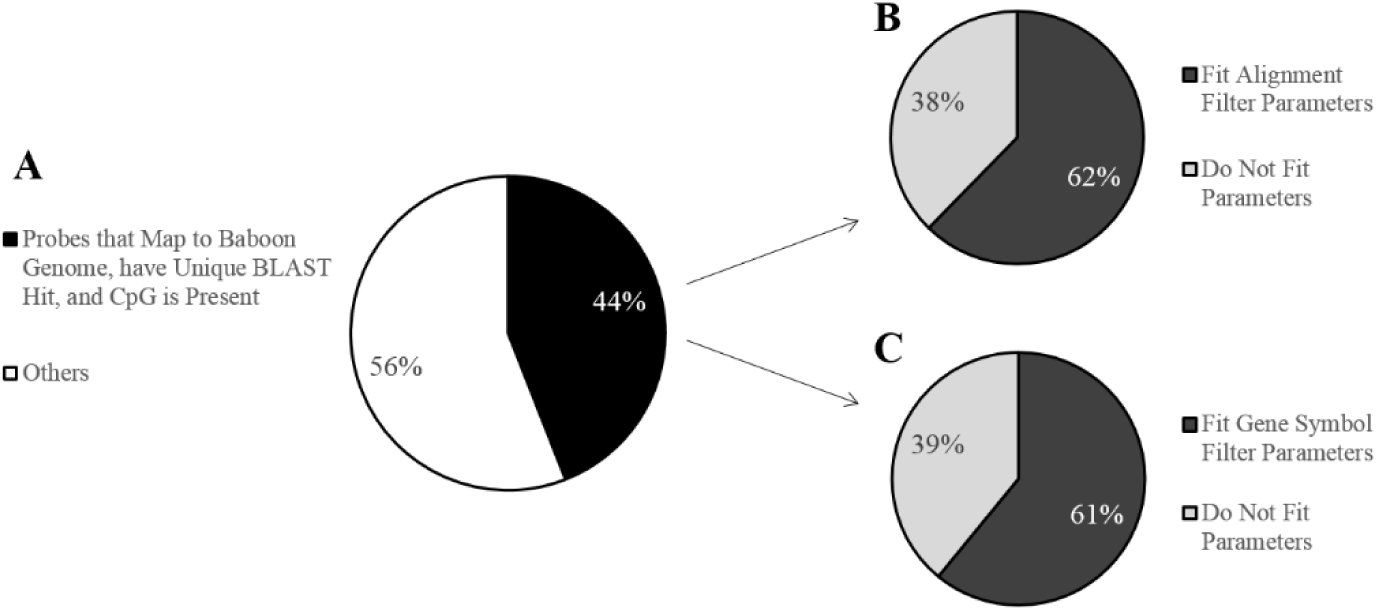
Filtering effects on the 450K array probes for baboons. **A,** Pie chart showing the percent of 450K array probes that map to the baboon (*Papio anubis*) genome with e-values less than e^-10^, have only unique BLAST hits, and target a CpG site. Out of 485,512 probes total, 213,858 probes (44%) meet these criteria. **B,** Pie chart showing the percent of probes, out of those that successfully mapped to the baboon genome, that contain 0 mismatches in 5bp of the probe by and including the targeted CpG site and 0-2 mismatches in 45bp of the probe not including the CpG site. Out of the 213,858 mapped probes, 133,264 probes (62%) meet these criteria. **C,** Pie chart showing the percent of probes, out of those that successfully mapped to the baboon genome, with gene symbol matches to humans. Out of the 213,858 mapped probes, 130,307 probes (61%) meet these criteria.

### Effectiveness of450K Array Probes using Baboon DNA

To determine how effectively the 450K array probes measured DNA methylation in baboon DNA, we performed Spearman correlation tests between the hybridization efficiency of each probe and parameters defining the alignment quality of each probe to the baboon genome. Specifically, both probe alignment bitscores and percent identity were significantly negatively correlated with probe hybridization efficiency, and probe alignment e-values were significantly positively correlated with probe hybridization efficiency, regardless of filtering criteria (Table S3). However, filtering probes using the alignment filter criteria retained proportionally more successfully hybridized probes than filtering probes using the gene symbol filter criteria (Figure S3). Thus, filtering probes using the alignment filter criteria likely produces more reliable results.

### Differential Methylation and Osteoarthritis

Significant DMPs were only identified between healthy and OA individuals in cartilage tissues (Table S4). All of these DMPs displayed decreased methylation in OA cartilage samples as compared to healthy cartilage samples, and some of these patterns overlapped with those previously identified in humans. Conversely, no DMPs were found between tissue types or between disease states in bone tissues. When filtering probes using the alignment filer criteria, six significant DMPs were identified between OA cartilage samples and healthy cartilage samples, while only two DMPs were identified when filtering probes using the gene symbol criteria (Table 1). One locus, *RFXAP*, matched between these filtering methods. *RUNX1* has previously been found to be differentially methylated in OA and healthy cartilage in humans, with OA cartilage having lower methylation as compared to healthy cartilage.^5^ The other genes associated with these probes (*KLHL26*, *RFXAP*, *MIR497*, *MIR195*, *ELF1*, *ACSL1*, and *CMIP*) have not previously been associated with OA-related differential methylation in humans.^4–10^

**Table 1.** Significant DMPs Between OA Cartilage and Healthy Cartilage.^a^

| | Probe ID | $\Delta M$ | P-Value | Human Gene Symbol | Human CpG Position | Human Gene Group | Human CpG Island | Relation to CpG Island | Other Human OA Genes | Baboon Gene Symbol | Baboon CpG Position |
| --- | --- | --- | --- | --- | --- | --- | --- | --- | --- | --- | --- |
| Alignment Filter Probes | cg05295841 | -3.6136 | 0.0247 | *KLHL26* | chr19:18771126 | Body | chr19:18771078-18771340 | Island | *CRTC1*; *CRLF1*; *COMP* | *CRTC1* | chr19:17216922 |
|  | cg17890983 | -1.9683 | 0.0247 | *RFXAP* | chr13:37394336 | Body | chr13:37393213-37394092 | South Shore | *SMAD9* | *RFXAP* | chr17:15599705 |
|  | cg02329670 | -1.4981 | 0.0411 | *MIR497*; *MIR195* | chr17:6921400 | TSS200; TSS1500 | chr17:6925400-6927527 | North Shelf | *ALOX12* | *LOC103878622* | chr16:6672282 |
|  | cg18456803 | -2.0478 | 0.0411 | *ELF1* | chr13:41593519 | TSS200 | NA | NA | NA | *WBP4*; *LOC103879193* | chr17:19630529 |
|  | cg13030790 | -3.5437 | 0.0411 | *RUNX1* | chr21:36421503 | 5'UTR; 1stExon | NA | NA | NA | *RUNX1* | chr3:11500594 |
|  | cg24721647 | -2.1838 | 0.0411 | *ACSL1* | chr4:185726836 | 5'UTR | chr4:185724434-185724647 | South Shelf | NA | *ACSL1* | chr5:173909497 |
| Gene Symbol Filter Probes | cg17890983 | -1.9679 | 0.0374 | *RFXAP* | chr13:37394336 | Body | chr13:37393213-37394092 | South Shore | *SMAD9* | *RFXAP* | chr17:15599705 |
|  | cg04759112 | -1.6320 | 0.0381 | *CMIP* | chr16:81663392 | Body; Body | NA | NA | NA | *CMIP* | chr20:63369656 |

## Discussion

Here we used the 450K array to identify DNA methylation variation in bone and cartilage tissues from a baboon model of OA. This was done both to determine the effectiveness of this application for baboon DNA and to assess the evolutionary conservation of epigenetic-OA associations in the primate lineage. We show that using the 450K array is feasible in baboon tissues. *In silico* probe filtering methods^14,15^ indicated that 44% of all human probes could be reliably mapped to the baboon genome and contained an informative CpG. This number was lower than expected since previous researchers were able to reliably use these same methods to map 61% of the human probes to the Cynomologus macaque genome,^15^ another Old World monkey that is a close phylogenetic relative to baboons. This discrepancy in number may be due to the quality of each nonhuman primates’ genome assembly. While both are well annotated, the average scaffold length (88,649,475) and contig length (86,040) of the macaque genome (Assembly: Macaca_fascicularis_5.0, Accession: GCF_000264685.2) are higher than those (528,927 and 40,262, respectively) of the baboon genome.

Subsequent *in silico* analyses based on sequence alignment criteria^14^ and based on gene symbol criteria^15^ retained similar numbers of probes (Figure 3) that maintained wide and comparable distributions throughout the genome (Table S2). However, only a little more than half of the resulting probes for each filtering technique overlapped with one another (Figure S2). This discrepancy is likely due to the incomplete nature of the baboon genome annotation. More than half of the probes that fit the alignment filter criteria but not the gene symbol criteria (28,699 out of 50,117) are associated with generic gene symbol identifiers (LOC) to indicate the as of yet unknown functions of these regions. Conversely, all of the probes that fit the gene symbol criteria but not the alignment filter criteria have over three mismatches with the baboon genome on average and have a maximum of nine mismatches with the baboon genome. These high mismatch numbers are a potential concern for proper and accurate probe and baboon DNA hybridization.

Fittingly, after applying the 450K array to measure DNA methylation patterns of genomic material extracted from baboon skeletal tissues, we found that the hybridization efficiency of probes was significantly correlated with the alignment quality of each probe to the baboon genome, and thus, the degree of sequence conservation. The majority of filtered probes for both *in silico* methods passed quality controls and produced robust signals on the array, indicating that either filtering technique may be appropriate for future research. However, because the filtering method based on the alignment filter criteria retained a larger proportion of successfully hybridized probes than the method based on the gene symbol criteria (Figure S3) and because this method is less influenced by the degree of genome assembly annotation, we recommend that this alignment filter criteria method be preferentially used in subsequent nonhuman primate studies.

This work is an extension of previous work using the 450K array to study DNA methylation patterns in the tissues of nonhuman primates.^14,15^ The 450K array is advantageous because it is cost efficient per sample and simultaneously measures a large number of CpG loci with a broad genomic representation.^38^ Similar to this study, previous researchers have applied the 450K array to measure DNA methylation patterns in great apes,^14^ which are closer to humans evolutionarily than baboons, and in macaques,^15^ which are comparable in proximity to humans evolutionarily as compared to baboons. In this study, we used a baboon model of OA to assess the evolutionary conservation of epigenetic-OA associations in the primate lineage, and we identified significant DMPs between healthy and OA individuals in cartilage tissues. All of these loci showed hypomethylation in OA cartilage samples as compared to healthy cartilage samples. Six DMPs were identified when using the alignment filter criteria, and two DMPs were identified when using the gene symbol filter criteria. All together, these eight DMPs are annotated to eight genes that have a variety of functions and are also in close proximity to five other genes that have previously been associated with OA (Table 1).

Some of these annotated genes have functions related to skeletal development and maintenance. For instance, *RUNX1* (Gene ID: 861), also known as runt related transcription factor 1, is involved in the regulation of bone and cartilage cell development and differentiation.^39^ Additionally, *MIR497*(Gene ID: 574456) and *MIR195* (Gene ID: 406971) are non-coding microRNAs that are involved in post-transcriptional regulation.^40^ While both of these microRNAs have roles in the development of cancer,^41,42^ they also play important regulatory roles in the differentiation of mesenchymal stromal/stem cells into bone related cells.^43^

Other genes have functions related to the immune system which may have proximal roles in the development of OA. In particular, *RFXAP* (Gene ID: 5994), also known as regulatory factor X associated protein, codes for a protein that assists in the transcriptional activation of major histocompatibility class II genes which are critical for the development and control of the immune system.^44^ Additionally, *CMIP* (Gene ID: 80790), also known as c-Maf inducing protein, codes for a protein that is involved in the T-cell signaling pathway, and SNPs within this gene have been associated with chronic diseases like diabetes.^45^

The remaining genes do not have functions related to skeletal phenotypes which makes their involvement in OA less clear. For example, *KLHL26* (Gene ID: 55295), also known as kelch like family member 26, is part of a family of proteins that may be involved in protein ubiquitination^46^. Additionally, *ACSL1* (Gene ID: 2180), also known as acyl-CoA synthetase long-chain family member 1, codes for a protein that assists in the biosynthesis of lipids and degradation of fatty acids, and SNPs within this gene have been associated with chronic diseases like diabetes.^47^ Lastly, *ELF1* (Gene ID: 1997), also known as E74 like E26 transformation-specific related transcription factor 1, is an important positive regulator of the Hox cofactor Myeloid ectropic viral integration site 1 (*MEIS1*) which is involved in developmental processes.^48^.

Out of all of these DMPs and their annotated genes, *RUNX1* is the only gene in which differential methylation has previously been associated with OA in humans. Specifically, *RUNX1* was found to be differentially methylated in OA and healthy cartilage in humans, with OA cartilage displaying hypomethylation as compared to healthy cartilage.^5^ Additionally, a drug targeting this gene has been proposed as a therapy for the treatment of OA.^49^ As of yet, none of the remaining DMPs and their annotated genes have been identified as candidate methylation loci in human OA studies.^4–10^ Nevertheless, some DMPs are located in close proximity to genes that have been previously associated with OA, including *CRTC1*^50^, *CRLF1*^51^, *COMP*^52^, *SMAD9*^53,54^, and *ALOX12*^55,56^ (Table 1). However, the mechanisms by which these DMPs might influence the expression of these distally related genes is unclear.

Overall, these findings suggest that one DNA methylation pattern associated with OA is evolutionarily conserved between humans and baboons. However, further work is necessary to fully elucidate whether the mechanisms contributing to *RUNX1* hypomethylation and its downstream effects in primate OA are also conserved, as well. Simultaneously, several DNA methylation patterns associated with OA do not appear to be evolutionarily conserved between humans and baboons. Differences between these two species may be due to general speciation events that took place during the evolution of these taxonomic groups, to slight differences in the development or manifestation of OA in these species, or artifacts of the experimental design itself. For instance, the sample size of this study (n=10) is rather small, and all individuals included were female. The small number of individuals likely reduced our power to detect potentially important OA-related DMPs, and the inclusion of only one sex may have biased our results such that identified OA DMPs are actually female specific DMPs. Thus, in order to improve the identification of candidate epigenetic alterations that underlie variation in knee OA, a larger sample set that includes both sexes should be considered. Nevertheless, using baboons as a model of OA in this study has begun to clarify the evolutionary conservation of this disorder, and future research in this animal model will help provide insight into the development and progression of OA in order to begin designing preventative and therapeutic agents.^16^

In conclusion, we determined that the 450K array can be used to measure genome-wide DNA methylation in baboon tissues and identify significant associations with complex traits. This is the first study to specifically assess DNA methylation in skeletal tissues from a nonhuman primate using this method. Some methylation variation is related to genes that impact skeletal development and maintenance, and this may have direct downstream regulatory and phenotypic effects. Additionally, while some DNA methylation patterns associated with OA in baboons appear to be evolutionarily conserved with humans, others do not. These findings warrant further investigation in a larger and more phylogenetically diverse sample set. Lastly, the work presented here begins to advance areas of research that incorporate an animal model of disease and an evolutionary perspective of diseases across phylogenies.

## Acknowledgements

We thank members of the Department of Genetics at the Texas Biomedical Research Institute for helpful discussions, including Erin Sybouts, Sophia Johnson, Mel Carless, Kara Peterson, Laura Cox, Kenneth Lange, Jerry Glenn, & Clint Christensen.

## Author Contributions

All authors were involved in drafting or critical revision of this article, and all authors approved the final version to be published. GH had full access to the study data and takes responsibility for the integrity of these data and the accuracy of their analysis. Study conception and design: GH, LMH, EQ, ACS.

Acquisition of data: GH, LMH, AGC.

Analysis and interpretation of data: GH.

## Declaration of Conflicting Interests

The authors of this manuscript declare that there are no conflicts of interest.

